# Performance modulations phase-locked to action depend on internal state

**DOI:** 10.1101/2022.11.28.518242

**Authors:** Tommaso Tosato, Gustavo Rohenkohl, Pascal Fries

## Abstract

Several studies have probed perceptual performance at different times after a self-paced motor action and found frequency-specific modulations of perceptual performance phase-locked to the action. Such action-related modulation has been reported for various frequencies and modulation strengths. In an attempt to establish a basic effect at the population level, we had a relatively large number of participants (n=50) perform a self-paced button press followed by a detection task at threshold, and we applied both fixed- and random-effects tests. The combined data of all trials and participants surprisingly did not show any significant action-related modulation. However, based on previous studies, we explored the possibility that such modulation depends on the participant’s internal state. Indeed, when we split trials based on performance in neighboring trials, then trials in periods of low performance showed an action-related modulation at ≈17 Hz. When we split trials based on the performance in the preceding trial, we found that trials following a “miss” showed an action-related modulation at ≈17 Hz. Finally, when we split participants based on their false-alarm rate, we found that participants with no false alarms showed an action-related modulation at ≈17 Hz. All these effects were significant in random-effects tests, supporting an inference on the population. Together, these findings indicate that action-related modulations are not always detectable. However, the results suggest that specific internal states such as lower attentional engagement and/or higher decision criterion are characterized by a modulation in the beta-frequency range.

## Introduction

The brain entails many rhythms, which in turn entail excitability fluctuations of the participating neurons. This likely influences the respective neuronal processing rhythmically, such that processing is modulated by the phase of the rhythm. Since behavior depends on neuronal processing, these rhythms can become directly visible in behavior. This can be investigated in at least two ways. First, by recording brain activity and showing that the phase of a rhythm modulates detection or discrimination accuracy (Busch et al., 2009). Second, by resetting a brain rhythm and showing frequency-specific modulations of task performance phase-locked to the reset. As a reset, an external event can be used, such as a flash or a sudden sound, which probably acts by resetting the phase of some internal rhythms in a bottom up fashion (Fiebelkorn et al., 2013; Landau and Fries, 2012). Alternatively a self-paced motor action can be used, such as an arm movement, a saccade, or a button press (Benedetto and Morrone, 2019; Benedetto et al., 2016; Tomassini et al., 2015). This may also act by resetting the phase of some internal rhythms, either by a corollary discharge signal sent from motor to sensory areas, or by sensory re-afference. Additionally, a motor action may reveal the phase of some internal rhythms modulating the likelihood of initiating that movement.

Recently, there was a debate about reproducibility of psychophysics studies reporting such modulations (Lin et al., 2021; Morrow and Samaha; van der Werf et al., 2021) and about the correct use of analysis methods in the field (Brookshire, 2022; Re et al., 2022; Tosato et al., 2022; Vinck et al., 2022). The fact that different studies applied different analysis methods and statistical tests makes the comparison difficult. Moreover, published studies show substantial discrepancy with regard to frequency and strength of effects. For example theta- or alpha-band centered modulations have been reported in different studies (VanRullen, 2016). Some of the variability has been addressed as a consequence of differences with regard to task difficulty (Chen et al., 2017), background luminosity (Benedetto et al., 2016), and attention (Busch and VanRullen, 2010; Harris et al., 2018).

Intriguingly, spontaneous slow fluctuations between brain states characterized by different oscillatory signatures have been described (Lakatos et al., 2016). This might well lead to different rhythms transpiring into behavior, within one participant at different times (assuming that the different states are alternating during the experimental session), or across different participants with different biases to one or the other state.

The debate about reproducibility might benefit from a study that uses a basic, simple paradigm in a relatively large sample size, and a statistical approach allowing an inference on the population. Therefore, we performed a study on 50 participants, using a self-paced button press followed by a detection task.

To our surprise, averaging our data over all trials of all participants, we found no significant frequency-specific modulation of perceptual performance phase-locked to the motor action. However, when we split trials based on performance in neighboring trials, we found an action-related modulation centered at ≈17 Hz. When we split trials based on the performance in the preceding trial, we found that trials following a “miss” showed an action-related modulation at ≈17 Hz. Finally, when we split participants based on their false-alarm rate, we found that participants with no false alarms showed an action-related modulation at ≈17 Hz. A 17 Hz behavioral modulation suggests an underlying neuronal beta rhythm.

## Results

Participants performed a self-paced button press, followed by a detection task on a probe that was presented at variable probe onset intervals (POIs), spanning from 0.1 to 1.1 s, after the button press (Figure 1). The probe’s contrast was adjusted with a staircase procedure such that performance was maintained close to 50%.

**Figure 1.**
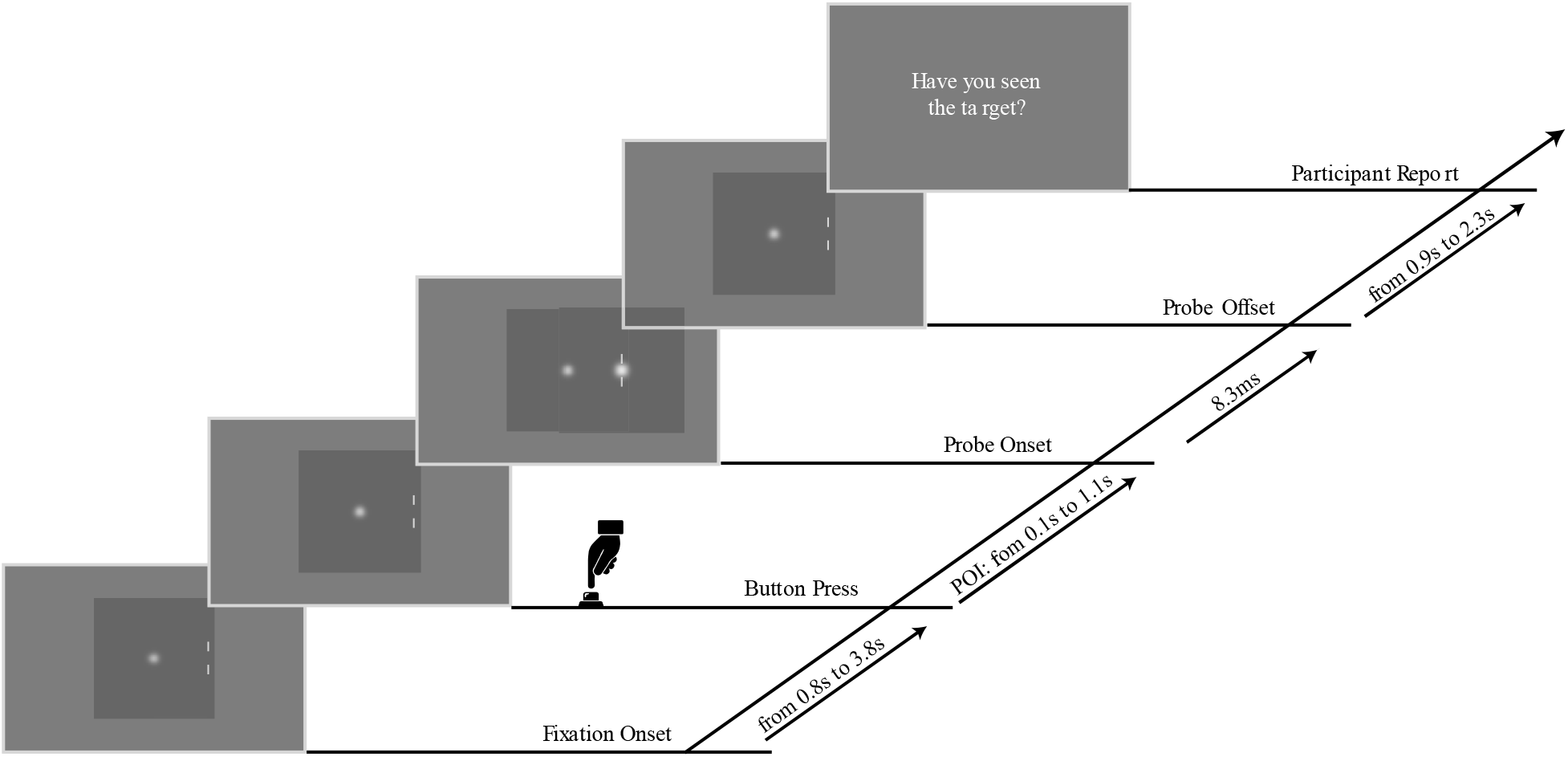
Task Paradigm. The trial started with the display of a fixation point and two fiducial lines on a gray background. Participants were instructed to perform a self-paced button press in a time window ranging from 0.8 to 3.8 s after the start of the trial. Following the button press, a probe appeared in 90% of the trials. Probe onset intervals (POIs) ranged from 0.1 to 1.1 s. After an additional delay, the question “Have you seen the target” appeared on the screen, and participants responded “Yes” or “No” on a keypad.

### Analysis of the aggregate data

Each trial provided a “hit” or a “miss”, referred to as behavioral response value (BRV). We analyzed all trials of all participants, testing for the hypothesized modulation of accuracy that was phase-locked to the reset event, here the button press (Tosato et al., 2022). We calculated an accuracy time course (ATC) for each participant by convolving the BRVs, which were aligned accordingly to their POIs, with a Gaussian kernel. Each ATC was then linearly detrended, and the detrended ATCs of all participants were averaged to give the mean ATC, shown in Fig. 2A. The phase-locking spectrum was estimated by means of a single-trial least square spectral analysis (stLSSA) applied to the trials of each participant separately, after linear detrending and Hann tapering (Tosato et al., 2022). In this way we obtained a complex spectrum per participant, and we then averaged the spectra of all participants. Lastly, we took the absolute values of the average spectrum, and we squared it to obtain the power.

**Figure 2.**
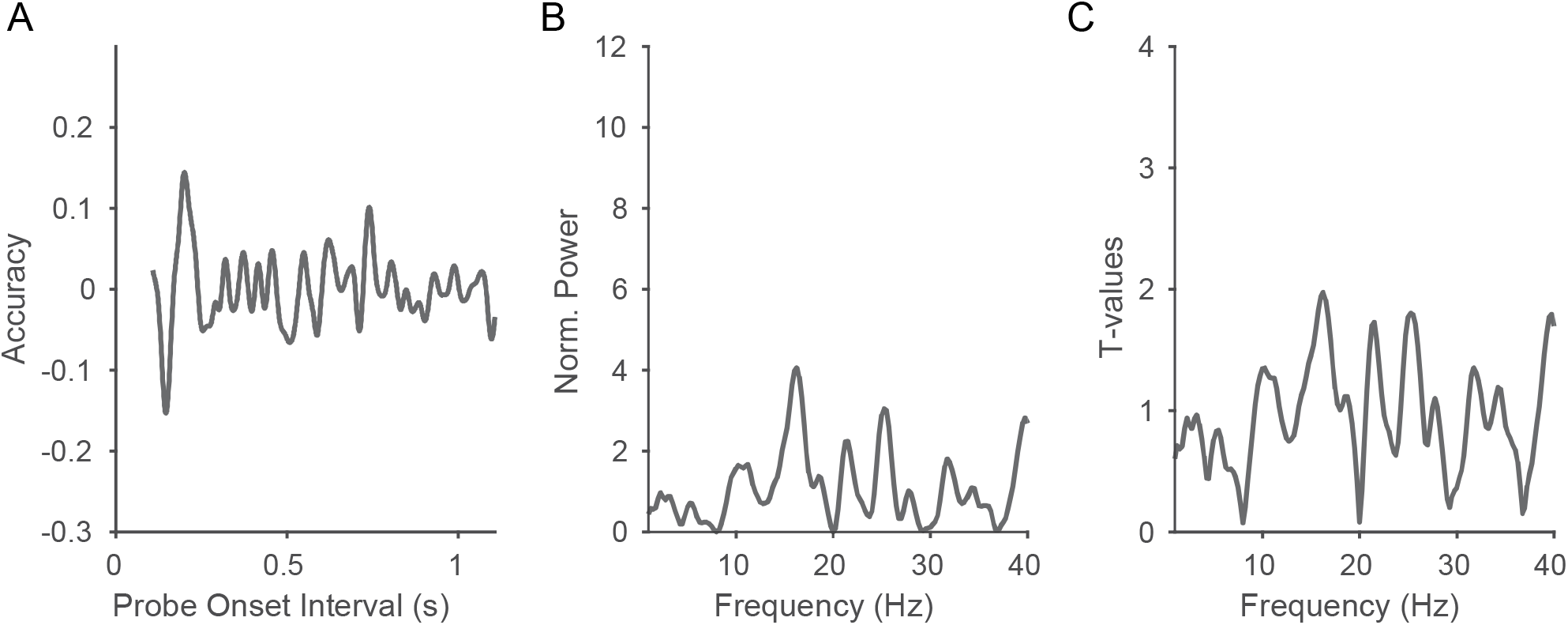
Spectral analysis of the pooled data. A) Mean accuracy time course obtained by averaging the linearly detrended accuracy time courses of each participant. B) Power spectrum obtained by squaring the absolute values of the average over the individual participants’ spectra. C) T-value spectrum resulting from paired t-tests comparing the observed spectra and the bias-estimate spectra across all participants

To determine significance, we first performed a fixed-effects non-parametric statistical test. To do so, we compared the values of the spectrum of the observed data to a distribution of values obtained by shuffling 5000 times the POIs of each participant and repeating the same analysis performed for the observed data. The observed spectrum was normalized by the mean of the spectra of the permuted data, for a better comparison with the following analysis. No frequency bin reached significance (p<0.5) after a max-based multiple comparison correction (Fig. 2B).

We then performed a random-effects statistical test. To do so we calculated for each participant a bias-estimate spectrum (see Methods). We then performed a paired t-test to compare the observed spectra and the bias-estimate spectra across participants, thereby obtaining the observed t-value spectrum (Fig. 2C). These t-values where then compared to a distribution of t values obtained after randomly exchanging in each participant the observed spectra and the bias-estimate spectra. No frequency bin reached significance (p<0.5) after a max-based multiple comparison correction.

### Task epochs with lower average performance show ≈17Hz accuracy modulation

It is known that the brain switches between different states over time (McGinley et al., 2015). For example, states of relatively strong theta and weak alpha rhythmicity in sensory areas are a signature of strong attentional engagement (Lakatos et al., 2016). Moreover it is known that participants do not sustain a constant level of attention during a task, but they repeatedly drift into mind wandering (Mooneyham and Schooler, 2013). We hypothesized that our participants may have gone through different states, for example of higher and lower attentional engagement. We reasoned that different levels of attentional engagement would have differently affected task performance. Therefore, we quantified performance for neighboring trials, referred to as local performance, and investigated effects of local performance on action-related modulation. Periods in the session where the local performance was higher than the overall performance were referred to as “*high-performance epochs”* and those where it was lower as “*low-performance epochs”* (see Methods). We split the trials based on whether they had occurred in high- or low-performance epochs, and repeated the above analyses for those two groups of trials (Fig. 3A-C). *Low-performance epochs* resulted in significant action-related modulation at ≈17 Hz, which was significant both for a fixed-effects (Fig 3B) and for a random-effects test (Fig 3C). For high-performance epochs, there was no such effect.

**Figure 3.**
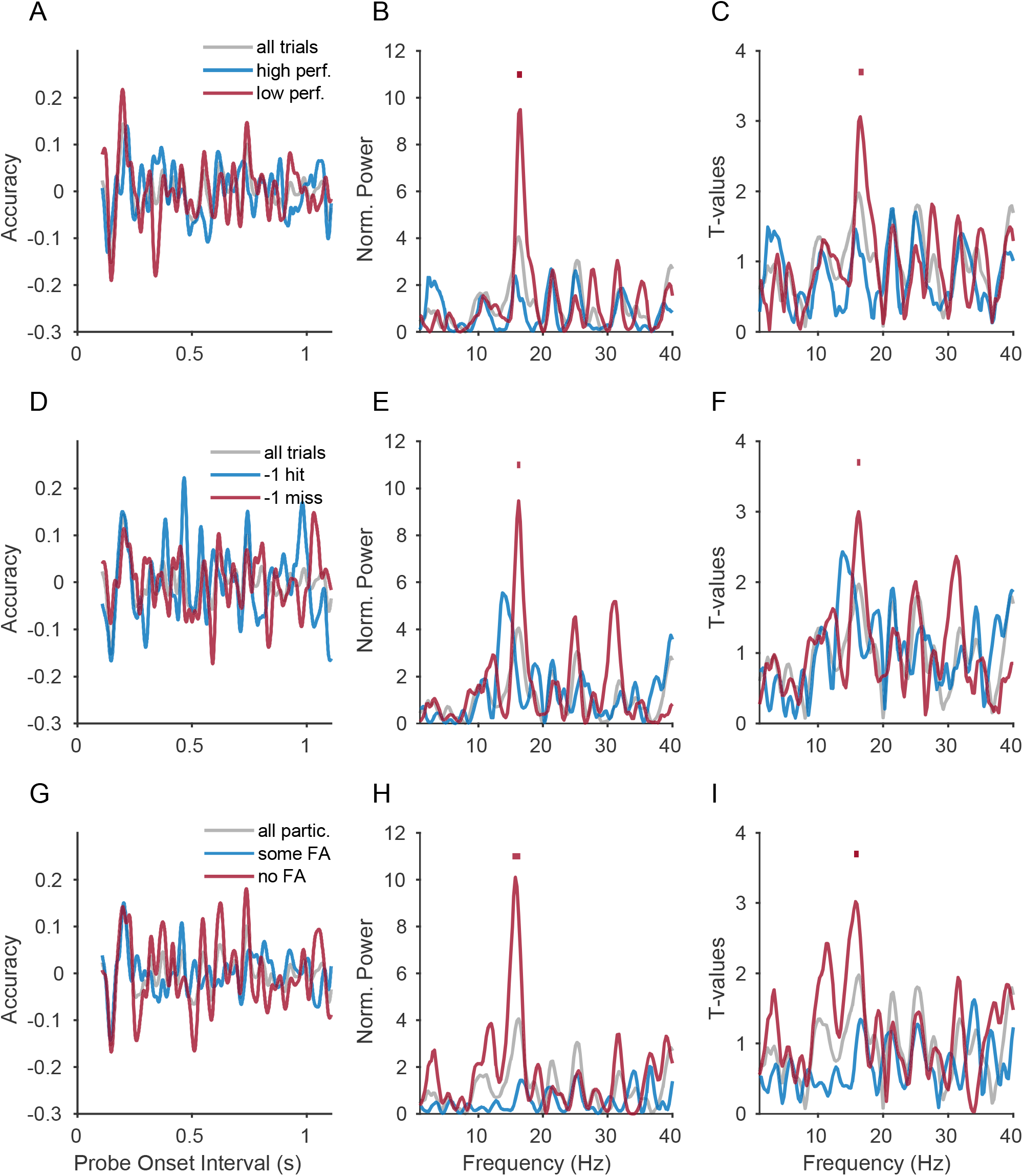
Spectral analysis of subsets of trials or participants. (A) Mean ATC, calculated separately for the trial group “*low performance epochs”* (red line), “*high performance epochs”* (blue line), and for all trials pooled (grey line). (B) Using the color code of (A), spectra show results for fixed-effects test, as described in Results and Methods. Significant frequency bins are marked above the spectra in the corresponding color. (C) Same as (B), but for random-effects test. (D-F) Same as A-C, but for the trial split indicated in D, namely “trial -1 miss” (red), “trial -1 hit” (blue), and all trials (grey). (G-I) Same as A-C, but for the participant-split indicated in G, namely “no FA” (red), “some FA” (blue), and all participants (grey)

In a separate analysis, we confirmed that low- compared to high-performance epochs had lower accuracy (Fig. 4A, p<0.001). Intriguingly, low-compared to high-performance epochs also had smaller pupil diameter (Fig. 4B, p<0.001) and higher microsaccade rate (Fig. 4C, p=0.003). Lower accuracy, smaller pupil diameter and higher microsaccade rate are consistent with states of lower attentional engagement (Bosman et al., 2009; Denison et al., 2019; Pastukhov and Braun, 2010).

**Figure 4.**
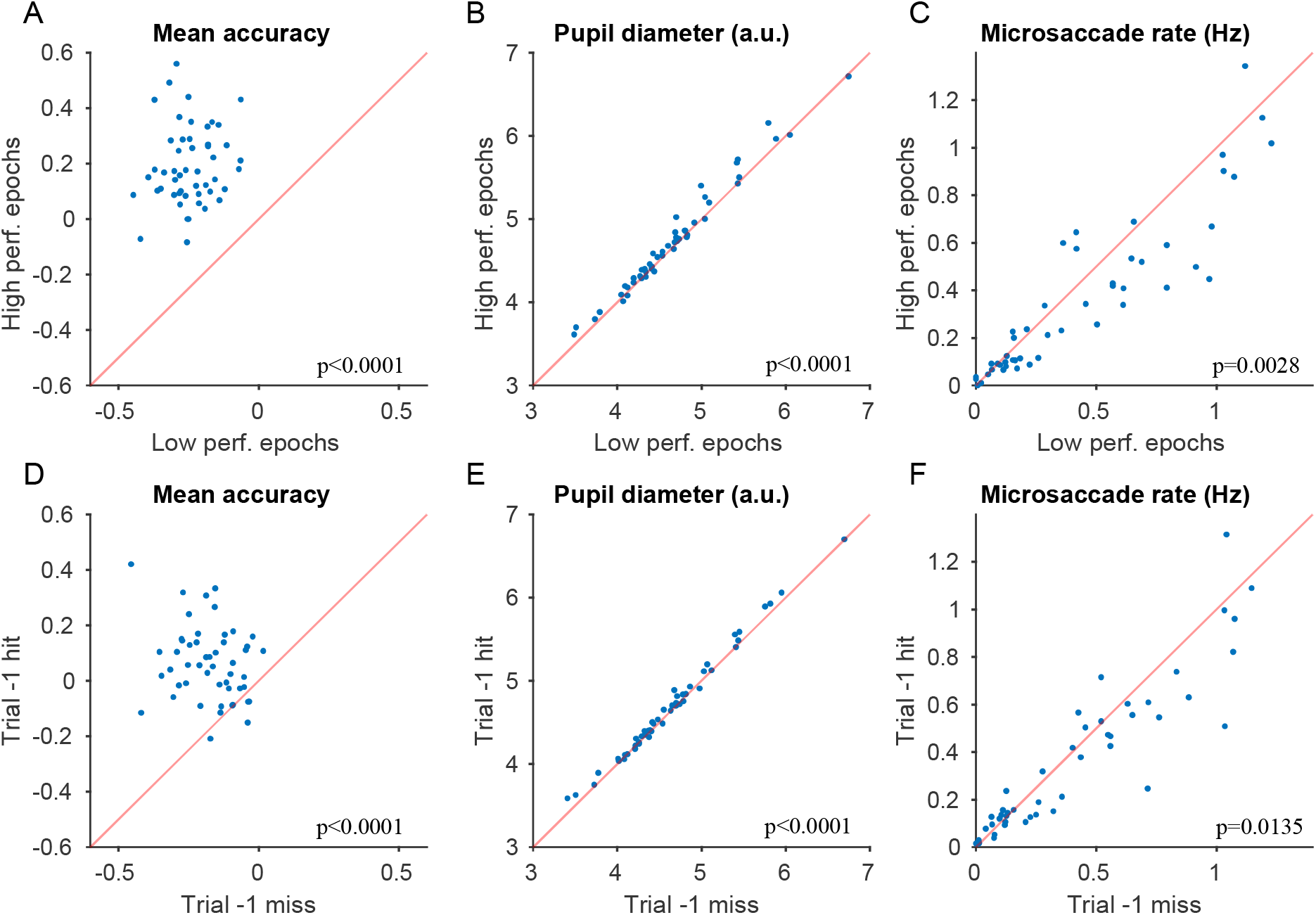
Characterization of subsets of trials. (A) Scatter plot of mean accuracy for trials from “*low performance epochs*” versus “*high performance epochs*”; each dot corresponds to one participant. (B) Same as A, but showing mean pupil diameter. (C) Same as A, but showing microsaccade rate. (D-F) Same as (A-C), but for “trial -1 miss” versus “trial -1 hit” trials.

### Trials following a missed detection show ≈17 Hz modulation

A recent study showed that the dynamic of rhythmic sampling in a given trial is influenced by the stimulus identity in the previous trial (Bell et al., 2020). The authors speculated that this is due to different predictions or expectations induced by the previous stimulus. Inspired by this study, we decided to split our trials based on the previous trial’s performance. We therefore divided our trials in two groups based on the performance history, and we referred to them as *“trial -1 hit”* and *“trial -1 miss”* (Fig. 3D-F; the first trial of each block was excluded as it had no immediately preceding trial). The *“trial -1 miss”* group showed significant action-related modulation at ≈17 Hz, which was significant both for a fixed-effects (Fig 3E) and for a random-effects statistical approach (Fig 3F). For the *“trial -1 hit”* group, there was no such effect.

In a separate analysis, we found that “trial -1 miss” compared to “trial -1 hit” trials had lower accuracy (Fig. 4D, p<0.001). Intriguingly, “trial -1 miss” compared to “trial -1 hit” trials also had smaller pupil diameter (Fig. 4E, p<0.001) and higher microsaccade rate (Fig. 4F, p=0.014). As mentioned above, this is consistent with states of lower attentional engagement.

### Participants with no false alarms show ≈17 Hz accuracy modulation

Finally, we explored the possibility that the number of false alarms (FAs) in each participant may correlate with the presence or absence of the ≈17 Hz modulation. We therefore split participants based on the total number of FAs (Fig. 3G-I). Participants who performed one or more false alarms constituted the group referred to as “*some FA”* (26 participants), and participants who did not perform any false alarms constituted the group referred to as “*no FA”* (24 participants). The “*no FA*” group showed an action-relation modulation at ≈17 Hz, which was significant both in a fixed-effects (Fig 3H) and in a random-effects (Fig 3I) statistical test. The “*some FA”* group showed no such effect.

## Discussion

In summary, when all participants and all trials were combined, the self-paced button press was not followed by frequency-specific modulations of perceptual performance phase-locked to the action. However, when participants or trials were split based on behavioral performance, action-related modulation emerged. When trials were split based on hit rate in neighboring trials, the subset of trials with low hit rate showed a ≈17 Hz modulation. Similarly, when trials were split based on whether the previous trial was a miss, the subset of trials following miss trials showed a lower hit rate and a ≈17 Hz modulation. For both of these trial splits, the trial subset with lower performance also showed smaller pupil size and higher microsaccade rate. Finally, when participants were split based on false-alarm rate, the subset of participants without false alarms showed a ≈17 Hz modulation.

Several seminal studies have reported periodic modulations of accuracy time courses aligned to a voluntary motor action, such as an arm movement (Tomassini et al., 2017; Tomassini et al., 2015), a button press (Benedetto et al., 2016; Nakayama and Motoyoshi, 2019; Zhang et al., 2019), or a saccade (Benedetto and Morrone, 2017; Hogendoorn, 2016; Wutz et al., 2016). Other studies have reported accuracy modulations after a sensory event, such as a visual flash (Landau and Fries, 2012; Re et al., 2019; Song et al., 2014) or an auditory input (Ho et al., 2017; Plöchl et al., 2021; Romei et al., 2012). These studies are based on the assumption that sensory events reset, and motor actions reset and/or reveal the phase of brain rhythms, allowing the experimenter to align trials to that event.

In contrast to our initial hypothesis based on these findings, we did not find any significant accuracy modulation after button press in our pooled data. This result adds to other recent reports of inconclusive and null findings (Morrow and Samaha, 2021; van der Werf et al., 2021; Vigué-Guix et al., 2020) and it is in line with the general revision happening in the field (Keitel et al., 2022). However, this result does not question the validity of the studies implementing similar paradigms (Benedetto et al., 2016; Nakayama and Motoyoshi, 2019; Zhang et al., 2019), because of differences in the task design. For example, we used a detection task with small stimuli presented in the periphery, instead of a discrimination task with larger stimuli presented in the fovea.

We explored the possibility that a modulation was present in a subset of trials and participants, by performing a post-hoc splitting of the data. This revealed significant phase locking at ≈17 Hz in trials occurring during task epochs characterized by lower mean performance, in trials occurring after a missed detection, and in participants that did not commit false alarms. The significant frequency bins at ≈17 Hz were part of a peak that stood out clearly in the spectrum, which is suggestive of an underlying beta-band rhythmicity. Note that spurious peaks in such phase-locking spectra can arise by a combination of higher-order detrending, amounting to high-pass filtering, with binning, amounting to low-pass filtering (Tosato et al., 2022). Therefore, we used neither higher-order detrending nor binning. Furthermore, when these data analysis steps produce spurious peaks, they are typically at the low end of the spectrum, whereas we found peaks consistently at ≈17 Hz.

Typical modulation frequencies in previous studies were lower in frequency, in the theta or alpha range. The finding of a modulation in the beta range however is in line with recent reports. Veniero et al. (2021) applied a TMS pulse on FEF while participants were performing a motion discrimination task, and immediately after the TMS pulse, task performance was modulated at ≈17 Hz. In the same study, the TMS pulse on FEF was shown to reset the phase of the low-beta oscillations recorded by the EEG occipital channels. A similar reset of beta oscillations may have happened in our experiment due to a corollary discharge in FEF, or related prefrontal/premotor region, linked to the button press. Other studies stressed the importance of beta oscillation for attentional processes in FEF (Fiebelkorn et al., 2018) and area V4 (Westerberg et al., 2021).

Bell et al. (2020) also reported a behavioral modulation in the beta range. Specifically they show a modulation of decision bias in a discrimination task of gender of androgynous faces, and the frequency of this modulation was either 14 Hz or 17 Hz depending on the identity of the previous stimulus. The fact that the previous stimulus influenced the accuracy modulation in the following trial is in line with our findings where the previous response influenced the modulation.

Other studies suggested that the frequency of rhythmic sampling may depend on different factors, such as task difficulty (Chen et al., 2017) and the attentional demands during the task (Merholz et al., 2022), which is in line with our interpretation that the action-related modulation might be related to attentional engagement. An additional explanation for the different frequencies reported in different studies is that different periodic modulations coexist in the brain, and the brain circuits recruited to perform a specific task, or reset by a specific event, may be under the influence of one or the other process (VanRullen, 2016).

We would like to offer two potential interpretations of the results of the analyses that splitted the data. The first interpretation pertains to the trial splits within participants and states that the presence of the ≈17 Hz modulation is a signature of lower attentional engagment in the task. Task epochs with lower performance may be a result of disengagement from the task, either because of fatigue, or mind wandering, or a combination of both. Trials following a missed detection were more likely to be a miss again, possibly because a state of lower attentional engagment was carried over from the previous trial. Visual attentional engagement in the absence of visual stimulation, as in the pre-probe period here, has been linked to reduced activity in the beta band, which includes the ≈17 Hz band found here (Siegel et al., 2008). The pupil and microsaccade data corroborate this interpretation, showing smaller pupil diameter and higher microsaccade rate in those trial subsets that contain the ≈17 Hz modulation. Small pupil diameter is known to be correlated with reduced arousal and locus coeruleus activation (McGinley et al., 2015).

An alternative interpretation pertains to the trial splits within participants and to the participant split, and it states that the presence of the ≈17 Hz modulation is a signature of a more conservative response criterion, i.e., a bias to report the absence of the target. Task epochs with lower performance may represent a higher criterion, leading to a higher proportion of misses. Trials following a miss were more likely to be a miss again, because a state of more conservative criterion was carried over from the previous trial. Participants with no false alarms performed the task with a higher criterion, which kept them from issuing false alarms. Beta band activity has been linked to mechanisms that maintain an existing state, or “status quo” (Engel and Fries, 2010; Gilbertson et al., 2005; Joundi et al., 2012), which might be related to a relative resistance against reporting a probe, which constitutes a visual change.

This study was not initially designed to perform the splitting procedures, and the task design was not adjusted for that purpose. However, the split over participants is orthogonal to the two splits over trials, and the fact that they produced very similar results provides some confidence in the validity of our findings. Here we recorded a larger sample size compared to previous experiments, and we performed both fixed-effects and random-effects statistical testing, using for the analysis the methods that, to our knowledge, provide the best trade-off between sensitivity and specificity (Tosato et al., 2022). We stress the need in the field for studies with higher statistical power (Button et al., 2013) random-effects statistics (Fries and Maris, 2021), and a careful consideration in the choice of the analysis methods (Kienitz et al., 2021; Tosato et al., 2022; Vinck et al., 2022).

In conclusion, when all trials were pooled, we did not find evidence for a consistent modulation of accuracy in a detection task where the probe was presented at variable times after a self-paced button press. However, we did find some indications for action-related modulation at ≈17Hz in a subset of trials and participants, which may have been characterized by lower attentional engagement or alternatively higher response criterion. These results should be taken with caution, because they were not initially hypothesized, and are the result of a post-hoc splitting. However they are suggestive of an interaction between attentional sampling dynamics and internal state, and further studies investigating this possibility may help to reconcile discrepancies between different studies in the field. Future experiments could therefore consider to mesure physiological indicators of arousal, such as pupil diameter, heart rate variability, respiration rate, skin conductance and subjective reports of absorption in the task (Whitmarsh et al., 2021).

## Methods

### Participants

Participants were recruited from the general public until 50 had successfully completed the task (25 women, average age 25.6). All had normal or corrected-to-normal vision and did not take any medication, except for contraceptives. All participants provided informed consent. The experimental protocol was approved by the ethics committee of the medical faculty of the Goethe University Frankfurt.

### Apparatus

The experiment was performed in a quiet, dimly lit room. Participants sat in front of a monitor (ViewPixx, diagonal: 57.15 cm; resolution: 1920 × 1200 px; refresh rate: 120 Hz) at a distance of 65 cm, with their head stabilized by a chin rest. Stimuli were generated in Matlab using Psychtoolbox-3. Eye position and pupil size were measured with an infrared eye tracker (EyeLink 1000, SR Research). Button presses at the beginning of each trial were recorded with a response box (fORP, 4 buttons Curved-Left, Cambridge Research Systems). The relevant time stamps were recorded by the EyeLink Host PC (both the ViewPixx monitor and the fORP response box were connected to the EyeLink screw-card, which was sampled at 1000 Hz by the EyeLink Host PC). Behavioral reports at the end of each trial were recorded with a numerical keypad.

### Stimuli and experimental procedure

The trial started with the presentation of a central fixation point (a 2 dimensional Gaussian of sigma = 0.5° of visual angle) on a gray background, and of two vertical fiducial lines situated around the horizontal meridian, 7.3° to the right of the fixation point. The participant was instructed to start fixating the fixation point as soon as it appeared and to keep fixating until it disappeared. Additionally, the participant was instructed to press a button on the response box, with the index finger of the right hand, in a time window ranging from 0.8 to 3.8 s after the start of the trial. The button press was followed by a probe onset interval (POI), ranging from 0.1 to 1.1 s, in which the participant had to monitor the location between the two fiducial lines for the appearance of the probe. The probe (a 2 dimensional Gaussian of sigma = 0.6°) was flashed for 8.3 ms (one frame given the monitor update frequency) between the 2 fiducial lines after the POI had elapsed. After a further variable time of 0.9 to 2.3 s, the fixation point and the fiducial lines disappeared from the screen, and the question “Have you seen the target?” appeared. The participant had to respond with the left hand pressing on a numeric keypad either the button 4 or 6, which were respectively labeled as “Yes” and “No”.

The POI assumed values ranging from 0.1 to 1.1 s, with a uniform probability distribution in time. 10% of the trials did not include a probe, i.e., were “catch trials”. Behavioral reports of probe detection were coded as 1 for a “Yes” response and -1 for a “No” response, and these values are referred to as behavioral response value (BRV).

Each participant performed a total of 430 trials distributed in 18 blocks of 24 trials each (∼140 seconds per block). After the completion of block 9, participants were asked to take a longer break and were offered water and a little snack. In 2 participants, this break occurred later (after block 11, and after block 14), and one participant skipped the break.

For each participant, a short preparatory session with a staircase procedure was used to obtain the participant-specific contrast threshold. The contrast resulting in 50% hit rate was selected as initial value for the main task. During the main task, the probe contrast was kept constant within each block. Between blocks, the contrast was slightly adjusted to maintain 50% hit rate and compensate for possible perceptual learning or tiredness.

The trials in which the button was pressed too early (earlier than 0.8 s after the start of the trial), or too late (later than 3.8 s after the start of the trial), and the trials in which no response was given, were marked as invalid and excluded from further analysis. Additionally, trials in which the gaze of both eyes was exceeding a distance of 50 pixels (corresponding to 1.1°) from the fixation point in the 100 ms preceding the probe onset were marked as invalid. 50 participants completed the task with 300 or more non-catch valid trials. 8 participants were excluded because they performed less than 300 non-catch valid trials.

### Accuracy time course

We calculated an accuracy time course (ATC) for each participant as a function of the POI time. We considered the time interval between 0.1 s and 1.1 s after the button press. The BRVs were arranged accordingly to their POIs, and they were convolved with a Gaussian of sigma=0.01 s in steps of 0.001 s. All the resulting time courses were linearly detrended and averaged over participants to give one mean ATC.

### Subdivision of the session into epochs

We subdivided each session into epochs characterized by higher or lower average performance. We calculated the time interval between the fixation onset of the first trial in the session and the probe onset of each trial, and called this measure “time in the session”. We analyzed task performance as a function of time in the session, by convolving the BRVs with a Hann window of 700 s length, and we referred to the obtained time course as local performance. The local-performance time course over the entire session was fit with a linear regression, to take into account a possible linear trend affecting performance along the session (e.g. decreasing performance due to increasing tiredness). Task epochs whose local performance was higher than the linear trend were labeled as “high performance epochs”; task epochs whose local performance was lower than the linear trend were labeled as “low performance epochs”.

### Microsaccades detection

Trials during which the eye signal deviated by more than 2° from the fixation point were excluded. The vertical and horizontal eye position signals (from 100 ms before to 1100 ms after the button press) were filtered with a low-pass Butterworth (2^nd^order with low-pass cutoff at 40 Hz). The first 100 ms were discarded to allow the filter to settle. The temporal derivatives of the filtered eye signals were taken as horizontal and vertical velocity, and they were combined to obtain overall eye speed. We used a speed threshold of 5 standard deviations (SD calculated per trial, across the entire 1100 ms, using a median estimation (Engbert and Kliegl, 2003)). Peaks in eye speed crossing the threshold where labeled as microsaccades. Only binocular microsaccades were used for further analysis. For each trial, we counted only the microsaccades occurring during the period when a probe could actually appear, i.e., between the button-press plus 100 ms and the appearance of the probe in that trial. The microsaccade rate of a participant (during a given condition, like low-performance epochs) was defined as the cumulative sum of microsaccade counts across trials (of that condition) divided by the cumulative time (in that condition), during which these microsaccades were detected.

### Spectral Analysis

To study phase locking effects at different frequencies, we applied the single-trial least square spectral analysis (stLSSA) (Tosato et al., 2022). As a preprocessing step, we linearly detrended and tapered the BRVs of each participant. To do so, we fit a line to the participant-specific ATC and we subtracted the value of this line at the time point of each trial’s POI from the respective BRV. Similarly, to taper single trials, the value of a Hann taper at the time point of each trial’s POI was multiplied with the respective BRV.

We then calculated a multivariate generalized linear model separately for each participant, using as independent variables, per frequency, the probe onset phases of all trials, and as dependent variables the corresponding BRVs. The model can be written as follow:

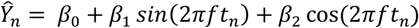

Where *Ŷ*_*n*_ are the predicted BRVs, *β*_0_ *β*_1_ *β*_2_ are the regression coefficients, *f* is the tested frequency, and *t*_*n*_ are the POIs. The regression parameters were estimated using the standard least square method as follow:

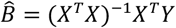

Where:

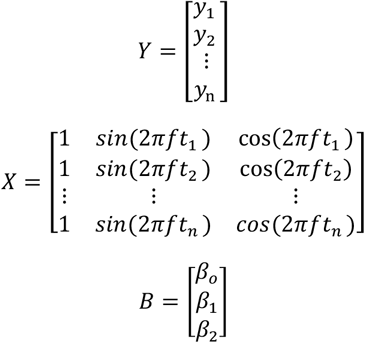

The regression coefficients *β*_1_ *β*_2_ were then combined as the real and the imaginary part of the complex Fourier coefficient for the frequency *f*. We repeated this procedure for all frequencies of interest, such that we had a complex number per frequency, which can be considered as the equivalent of a complex Fourier spectrum. As frequencies of interest, we used all frequencies between 1 Hz and 40 Hz, in steps of 0.25 Hz. These calculations were done at the level of the single participant, such that each participant gave one complex spectrum. The complex spectra of all participants were then averaged in the complex domain, i.e. taking phase information into account, to obtain a single complex spectrum. This complex spectrum was rectified and squared to obtain the power spectrum.

### Statistical Analysis

In this study we performed two different levels of inference: 1) An inference on the sample of investigated participants, referred to here as fixed-effects analysis. 2) An inference on the population of all possible participants (or all possible participants fulfilling a certain selection criterion, like no false alarms), referred to here as random-effects analysis.

The fixed-effects analysis was based on a randomization at the trials level. In this case we compared our observed spectra with a distribution of bias-estimate spectra. Each bias-estimate spectrum was obtained by randomly combining POIs and BRVs at the single participant level, calculating a spectrum per participant, and averaging those spectra over participants. This randomization was performed 5000 times, and from each resulting bias-estimate spectrum, we retained the maximal values across all frequencies. If the value for a given frequency in the observed spectrum was larger than the 95^th^ percentile of the maximal-value distribution, we considered that frequency bin significant at p<0.05 with multiple comparison correction across frequencies.

The random-effects analysis is based on a randomization at the participants level, and it allows us to make an inference at the population level. For each participant, we had the observed spectrum and the bias-estimate spectrum (averaged over the 5000 randomizations). A paired t-test between the observed and the bias-estimate spectra, across participants, separately per frequency, gave the observed t-value spectrum. We then randomly exchanged conditions at the level of the participant. That is: To implement one randomization, we make a random decision, per participant, of whether to exchange the observed spectrum with the bias-estimate spectrum, or not. We then proceed as before, arriving at one randomization t-value spectrum. This randomization was repeated 5000 times, and from each resulting t-value spectrum, we retained the maximal value across all frequencies. If the t-value for a given frequency in the observed spectrum was larger than the 95^th^ percentile of the maximal-value distribution, we considered that frequency bin significant at p<0.05 with multiple comparison correction across frequencies.

The testing procedures described in the last two paragraphs implement *multiple comparisons correction* across frequencies according to the max-based approach (Nichols and Holmes, 2002). After each randomization, the maximal value across all frequencies was placed into the maximal-value distribution. Note that those distributions lack a frequency dimension. The 95^th^ percentile of those distributions is the threshold for statistical significance with a false-positive rate of 0.05, including correction for the multiple comparisons across frequencies. Correspondingly, the observed power or t-value spectra were compared against those thresholds.

## Notes

### Competing Interest Statement

PF has a patent on thin-film electrodes (US20170181707A1) and is beneficiary of a respective license contract with Blackrock Microsystems LLC (Salt Lake City, UT), and he is member of the Advisory Board of CorTec GmbH (Freiburg, Germany). The authors declare that no further competing interests exist.

